# FliO is an evolutionarily conserved yet diversified core component of the bacterial flagellar type III secretion system

**DOI:** 10.1101/2025.05.06.652439

**Authors:** Ekaterina P. Andrianova, Amanda L. Dobbins, Marc Erhardt, David R. Hendrixson, Igor B. Zhulin

**Author notes:** Address correspondence to: Igor B. Zhulin or David Hendrixson. These authors contributed equally: Ekaterina P. Andrianova, Amanda L. Dobbins.

## Abstract

The bacterial flagellum is a complex nanomachine essential for motility, environmental sensing, and host colonization. While many of its core components have been well characterized, the relevance of proteins such as FliO, which are inconsistently annotated and poorly conserved at the sequence level, has remained ambiguous in their evolutionary and functional status. Here, we present a comprehensive phylogenomic and structural analysis of FliO across >30,000 representative genomes spanning >100 bacterial phyla. Additionally, during this analysis, we found that approximately 40% of bacterial genomes contain flagellar genes - significantly fewer than previously reported. Using a custom pipeline combining low-threshold HMM searches, operon context analysis, and structural information, we demonstrate that FliO is present in ∼95% of genomes encoding the core flagellar components FliP, FliQ, and FliR. This suggests that FliO is a nearly ubiquitous and ancestral core component of the flagellar type III secretion system (fT3SS). FliO exhibits considerable structural diversity, including lineage-specific acquisitions of LysM and AMIN domains. We identify FliO homologs not only in canonical flagellar systems but also in some virulence-associated T3SS and even some non-flagellated organisms, suggesting functional repurposing and highlighting its functional plasticity. Functional studies in *Campylobacter jejuni* reveal that FliO and its AMIN domain are critical for efficient bipolar flagellation, membrane stability of the export gate component FlhB, and colonization of the host. These findings establish FliO as a core, yet evolutionarily dynamic, component of flagella and provide new insights into the evolution and diversification of bacterial secretion systems.

## Introduction

Bacterial flagella are complex nanomachines that enable motility and are essential for a variety of cellular processes, including environmental sensing, surface attachment, and host colonization^1^. Flagella also serve as virulence factors in many pathogenic species, making them central to host-microbe interactions^2,3^. Despite advances in molecular and structural biology and comparative genomics, our understanding of flagellar evolution remains incomplete. In model organisms *Escherichia coli* and *Bacillus subtilis*, over 50 proteins are involved in flagellum assembly, regulation, and function^4,5^. Approximately half of these proteins are considered core components^6^, meaning they are evolutionarily conserved and present in all or nearly all flagellated species. This conservation across phylogenetically distant species suggests that the bacterial flagellum emerged early in evolutionary history and has been maintained through strong selective pressures due to its adaptive significance^6,7^. The other half of flagellar proteins exhibit a sporadic distribution across bacterial species, likely due to lineage-specific gene loss and acquisition. One of the proteins in this category is FliO, a component of the flagellar type III secretion system (fT3SS), a specialized export apparatus responsible for translocating structural flagellar proteins across the cytoplasmic membrane during assembly^2,8^. The fT3SS is evolutionarily and structurally related to the virulence-associated type III secretion system (vT3SS) found in many Gram-negative pathogens^2,8,9^. Both systems share a conserved architecture involving FliP, FliQ, FliR, FlhA, and FlhB – core components of the flagellar machinery that constitute the export gate complex^10^. The assembly of fT3SS begins with the formation of its core complex. In *E. coli* and *Salmonella enterica*, FliP is the first protein to integrate into the membrane, followed sequentially by FliR and FliQ to form a stable FliP₅Q₄R₁ secretion pore complex^10,11^. This complex is subsequently surrounded by a single FlhB subunit and a nonameric FlhA ring, forming the export gate of the fT3SS^12–14^.

In many bacteria, such as *E. coli, S. enterica,* and *B. subtilis*, FliO is encoded in an operon together with FliP, FliQ, and FliR. Previous studies suggested a functional link between FliO and FliP, supported by genetic evidence^15^. Subsequent research proposed that FliO acts as a flagellum-specific chaperone required for the efficient assembly of FliP-FliR oligomers^16^. In *S. enterica* and *Helicobacter pylori*, FliO orthologs are essential for flagellar biogenesis and motility^16,17^, Thus, it is surprising that FliO, in contrast to FliQ, FliP, and FliR, has not been universally recognized as a core component of the flagellar system.

Moreover, the functional role of FliO remains poorly understood. In *Salmonella*, FliO is thought to stabilize FliP and promote its integration into the membrane^16^. In *H. pylori*, FliO affects the stability of FlhA rather than FliP, likely because FlhA ring assembly requires the FliPQR core complex^17^, highlighting potential functional divergence across species. In contrast to the well-defined roles of other fT3SS components, FliO’s conservation, structural variation, and potential functional adaptability remain largely unexplored.

In this study, we developed a multi-tiered computational approach to identify FliO homologs across bacteria and performed a comprehensive phylogenomic and structural analysis to characterize the full diversity of FliO proteins. We show that FliO is nearly ubiquitous in flagellated bacteria, with rare losses and frequent structural innovations, particularly involving AMIN and LysM domains. To functionally validate these findings, we focused on FliO from *Campylobacter jejuni*, a flagellated intestinal commensal and human pathogen. Our experimental results demonstrate that FliO and its AMIN domain are critical for FlhB stability, efficient bipolar flagellation, and commensal colonization in the chick intestine. These findings establish FliO as a core component of the fT3SS and offer new insights into the processes driving the evolution of bacterial flagella.

## Results

### A computational strategy for identifying FliO homologs

The initial study reporting a sporadic distribution of FliO^6^ relied solely on BLAST^18^ as the similarity search tool. To assess this further, we performed standard BLAST searches using FliO sequences from two distantly related model organisms, *B. subtilis* and *E. coli* as queries against the same set of genomes. Strikingly, the *B. subtilis* FliO query failed to retrieve any hits, even in genomes of closely related species, such as *Listeria* and *Clostridium*, while the *E. coli* FliO query retrieved homologs only from closely related gammaproteobacteria (Supplementary Data 1). These failures are attributable to the high sequence divergence of FliO. For example, FliO sequences from *B. subtilis* (RefSeq accession NP_389516.1) and *E. coli* (NP_416457.4) share less than 30% identity over 76 aligned residues, yielding a BLAST E-value of 0.093, which is higher than the default significance threshold of 0.05. In contrast, FliP sequences from the same organisms (NP_389517.1 and NP_416458.1) are more than 50% identical over 208 aligned residues, with the highly significant E-value of 2e^-79^. These results indicate that BLAST frequently fails to detect FliO homologs, in contrast to other flagellar proteins, such as FliP.

PSI-BLAST^18^, a more sensitive profile-to-sequence search tool was previously employed to identify FliO homologs in *C. jejuni*^19^ and *H. pylori*^17^; however, in our hands, PSI-BLAST produced many spurious hits when applied to very large datasets, resulting in an unacceptably high rate of false positives. Fabiani *et al*^16^ employed a sequence-to-profile search tool, HMMER^20^, and Hidden Markov models (HMMs) from the Pfam database^21^ to scan 4,771 NCBI RefSeq genomes for the presence of FliO. They reported identifying FliO in approximately 80% of genomes that encoded other flagellar proteins, specifically FliP, FliQ, FliR, and flagellin. This finding supported a broader distribution of FliO than previously recognized and suggested its sporadic absence in several phyla. However, the underlying dataset was heavily biased - over 80% of genomes belonged to just four phyla - and no detailed genome-level results were provided. As a result, the true distribution of FliO across the >100 recognized bacterial phyla remains largely unknown.

To address this problem, we conducted a comprehensive analysis of the distribution of FliO proteins across 131 bacterial phyla, using a dataset of 30,238 representative genomes from GTDB database^22^. In parallel, we also analyzed for the presence of conserved core flagellar proteins, including components of fT3SS - FliP, FliQ, and FliR – as reference markers. We first employed KEGG HMMs to search for FliO, FliP, FliQ, and FliR across the dataset (see Methods). Using this approach, we identified FliP, FliQ and FliR in over 12,000 genomes, suggesting that approximately 40% of bacterial species are flagellated – noticeably lower than the previous estimate of 50%^23^. In comparison, FliO was detected in fewer than 11,000 genomes, corresponding to approximately 89% of genomes where FliP, FliQ, and FliR were found (Supplementary Data 2). Next, we repeated the search using the Pfam HMM for FliO, which yielded even fewer hits: only 76% of genomes containing FliP, FliQ, and FliR also had a detectable FliO (Supplementary Data 2).

The *fliO* gene is frequently located in the same operon as *fliP*, a relationship conserved across distantly related organisms, such as *E. coli* and *B. subtilis*. To investigate the possibility that current HMMs miss divergent FliO homologs, we examined the *fliP* gene neighborhoods in genomes where FliP, FliQ, and FliR were present, but FliO was not detected. In many such cases, we observed that genes whose products were annotated as “hypothetical proteins” were located adjacent to *fliP* within flagellar operons (Fig 1A). We subjected the protein products of these genes to sensitive profile-profile searches using HHPred^24^, which consistently identified them as FliO homologs with high confidence. These findings demonstrate that current HMMs do not detect many FliO homologs, likely due to stringent thresholds.

**Fig. 1.**
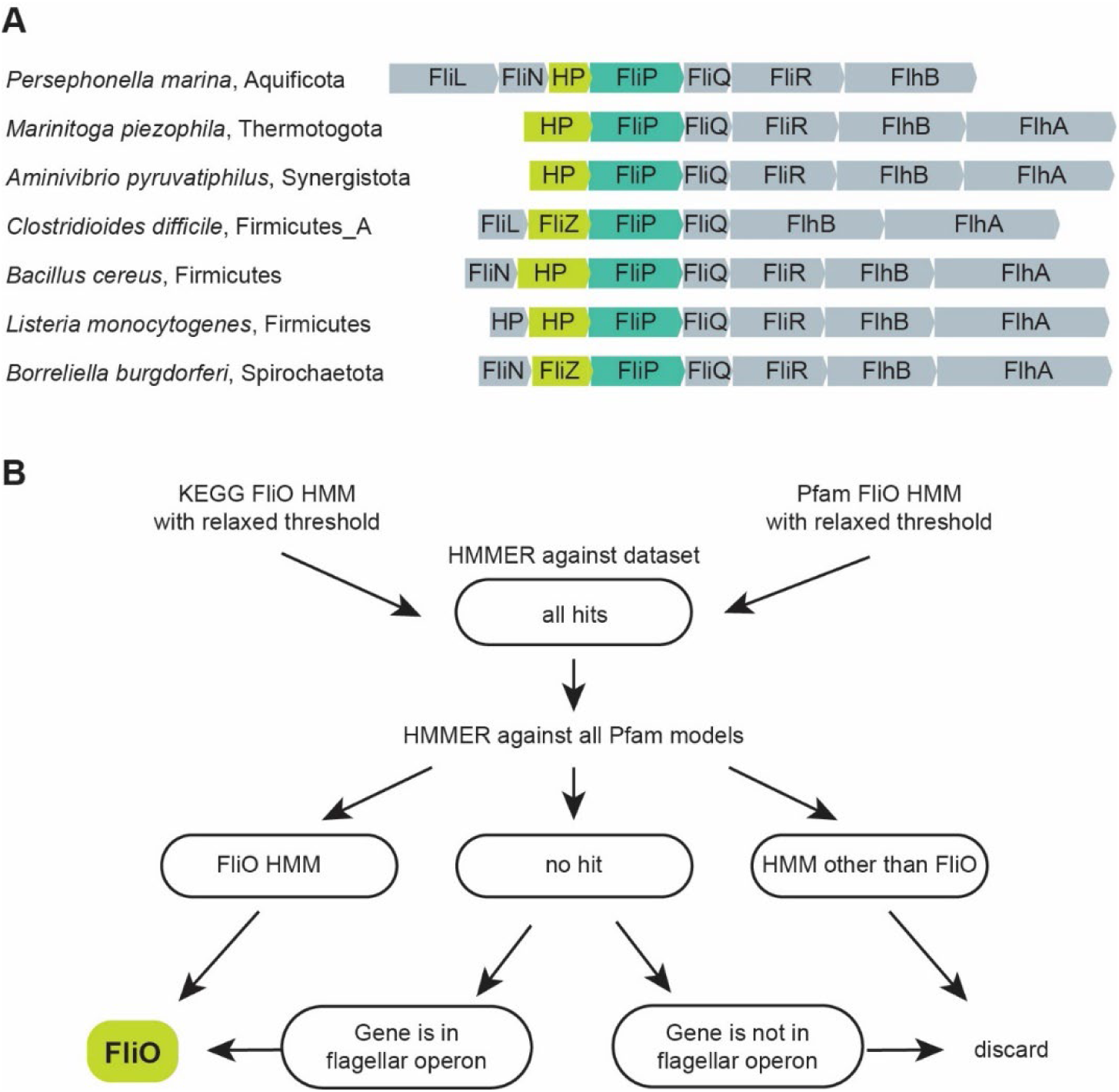
FliO poses a challenge for identification in genomic datasets using conventional bioinformatics approaches. (**A**) Representative examples of FliP gene neighborhoods, where FliO is annotated as a “hypothetical protein” (HP) or “FliZ” by bioinformatics tools implemented in the RefSeq database. FliP accession numbers: *P. marina*, WP_049756084.1; *M. piezophile* WP_014296225.1; *A. pyruvatiphilus*, WP_133955447.1; *C. difficile*, WP_003435977.1; *B. cereus*, WP_001220582.1; *L. monocytogenes*, WP_014600604.1; *B. burgdorferi*, WP_002556874.1. (**B**) Computational workflow for large-scale identification of FliO homologs in bacterial genomes.

To overcome the limited sensitivity of existing models, we conducted HMMER searches using both KEGG^25^ and Pfam HMMs against our genome dataset, applying a very low threshold to maximize the recovery of potential FliO homologs. While approach allowed us to detect numerous putative FliO sequences, it also substantially increased the rate of false positives. To address this, we implemented a multi-step filtering workflow to refine the results (Fig 1B).

First, we collected all sequences that matched either HMM at the low threshold. These sequences were then queried against the entire Pfam database using HHMER. Sequences that matched a different Pfam model with high confidence were discarded as non-relevant. Those that had a match to the FliO domain were kept and added to the curated FliO dataset. For sequences with no clear FliO domain prediction, we examined their gene neighborhoods. Proteins encoded by genes located adjacent to known flagellar genes were retained for further evaluation, while those found in non-flagellar neighborhoods and singleton genes were discarded. Using this strategy, we successfully identified FliO homologs in 11,705 genomes, representing 95% of the genomes that also contained FliP, FliQ, and FliR (Supplementary Data 2).

We then examined the distribution of additionally identified FliO homologs across bacterial phyla and observed that the detection failure rate varied significantly by lineage. In most phyla, FliO proteins were readily detected using default thresholds (Supplementary Data 2). However, in several phyla, a significant proportion of FliO homologs were only identified using the relaxed threshold. For instance, less than 45% of FliO proteins were detected using default search parameters in Synergistota, Aquificota, and Campylobacterota. Detection rates were also lower than expected in Thermotogota, Firmicutes, Spirochaetota, and Alphaproteobacteria (Supplementary Data 2).

### FliO homologs are present in all phyla of flagellated bacteria

We identified FliO homologs in 95% of genomes that also contain FliP, FliQ, and FliR. Analysis of the phyletic distribution of FliO homologs revealed that their consistent presence across all phyla of flagellated bacteria (Supplementary Data 2). This widespread distribution implies that FliO was already present in the last common ancestor of bacteria, which was suggested to be flagellated^26^, with subsequent losses occurring in only a few lineages. Notably, FliO appears to be absent due to gene loss in only two bacterial phyla: Campylobacterota and Pseudomonadota. In Campylobacterota, the absence of FliO can be attributed to a single evolutionary event, likely occurring in the common ancestor of the Arcobacteraceae family, where FliO cannot be identified in all 52 genomes containing FliPQR (Fig. 2, Supplementary Data 2), In contrast all other flagellated species within this phylum - including *H. pylori* and *C. jejuni* - retain at least one copy of the fliO gene. Interestingly, the flagellar operons in Campylobacterota exhibit a markedly divergent gene order compared to those in most other bacterial lineages (Fig. 2).

**Fig. 2.**
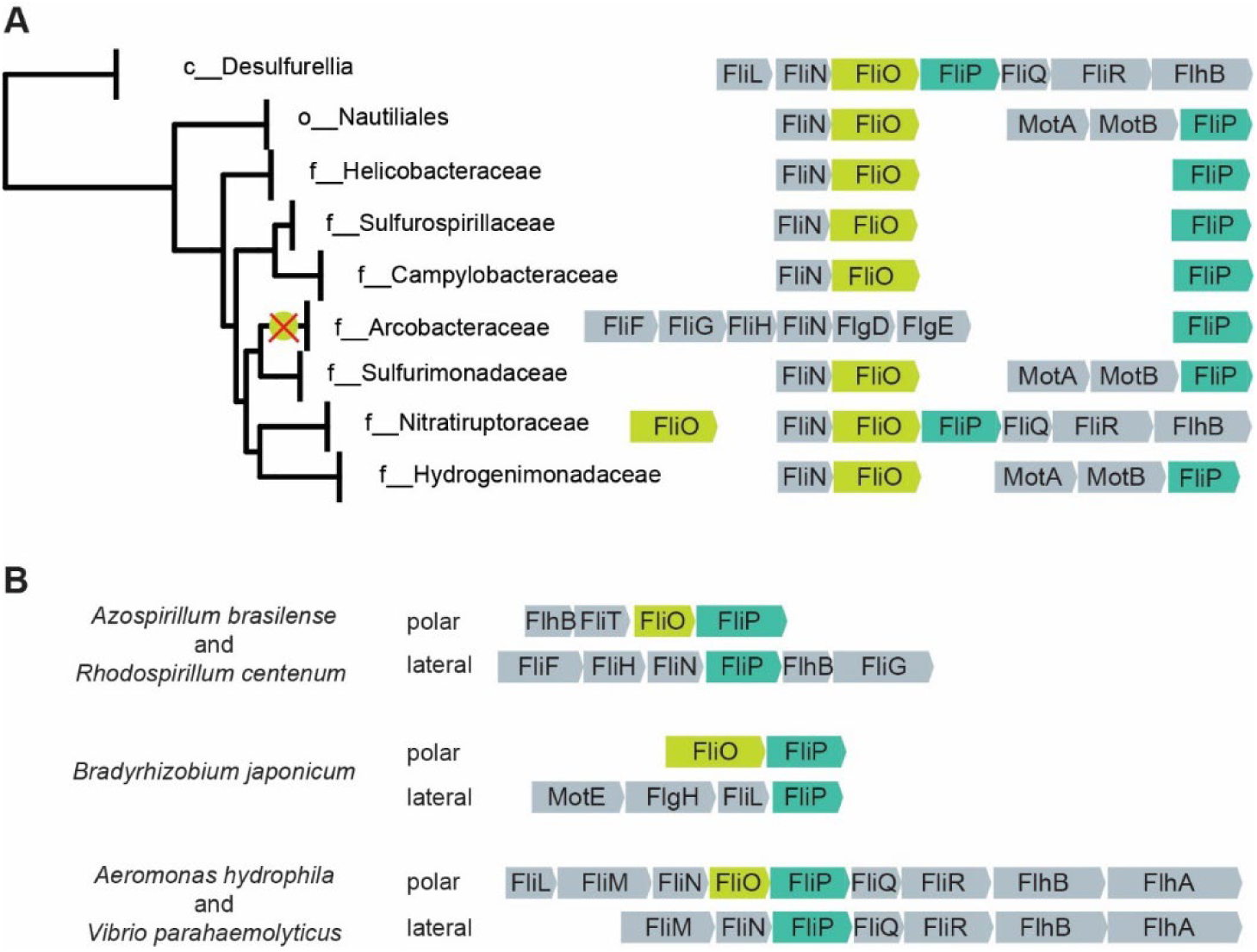
FliO gene loss in Campylobacterota and Pseudomonadota. (**A**) FliO loss in the family Arcobacteraceae (Campylobacterota). The loss of FliO is marked with an X. Diverse FliOP neighborhoods in the order Nautiliates are shown. The genome tree is based on GTDB database v95. (B) FliO loss in lateral, but not in polar flagella in representative species of Pseudomonadota.

In Pseudomonadota, the loss of FliO is more scattered. No FliO homologs were detected in various genomes belonging to alphaproteobacterial orders Azospirillales, Rhizobiales, Rhodobacterales, and Caulobacterales. Among Gammaproteobacteria, FliO was absent in a smaller subset of genomes, specifically within the orders Burkholderiales, Steroidobacterales, and Enterobacterales. Remarkably, in alphaproteobacteria, flagellar operons encoding FliO and FliP also exhibit substantial deviation from the canonical gene order (Fig 2).

Many species of alpha- and gammaproteobacteria possess two flagellar systems: a single polar flagellum and multiple lateral flagella. Each system is encoded by distinct sets of genes and is responsible for different types of motility^27^. We found that FliO is frequently absent from the set of flagellar genes required for the formation of lateral flagella and swarming motility, while it is present in polar flagellar systems responsible for swimming motility. For example, no FliO homolog was identified in the operons that encode lateral flagella in *Azospirillum brasilense, Rhodospirillum centenum*, *Bradyrhizobium japonicum*, *Aeromonas hydrophila*, and *Vibrio parahaemolyticus* (Fig 2).

### FliO in non-flagellated species and non-flagellar secretion systems

While the vast majority of FliO domain-containing proteins are encoded within flagellar operons, we also identified FliO homologs in non-flagellar contexts, indicating potential alternative functions. For example, non-flagellated Chlamydiia species from Verrucomicrobiota A phylum encode FliO homologs within a conserved five-gene operon of unknown function (e.g., WP_011097025.1) (Supplementary Data 2). Similarly, some members of the Rhizobiales order (Alphaproteobacteria) possess singleton FliO homologs - genes not located near any flagellar operons and found in genomes that lack other flagellar genes entirely (e.g., WP_132005128.1) (Supplementary Data 2). These findings suggest that FliO may have functions beyond the canonical flagellar system.

Strikingly, we also identified FliO homologs within vT3SS of certain bacterial species. Although FliO has traditionally been considered a flagellum-specific export apparatus protein and thought to be absent from non-flagellar T3SS^28^, our data show that FliO is indeed present in some of the non-flagellar T3SS operons (Supplementary Data 2). In the Myxococcota phylum, FliO is frequently encoded as a part of the vT3SS apparatus. For example, two distinct non-flagellar T3SS systems are present in *Myxococcus xanthus*, and both include a FliO homolog (WP_020477618.1 and WP_216608840.1). A recent study demonstrated that one of these systems is involved in initiating prey cell disintegration, which facilitates prey consumption, while the other vT3SS is not involved in prey killing^29^.

Additionally, we identified FliO in non-flagellar T3SS operons in several representatives of Pseudomonadota, including those from Caulobacterales, Sphingomonadales, Burkholderiales, Chromatiales, Granulosicoccales, and Xanthomonadales orders (Supplementary Data 2). In some of these genomes, *fliO* genes were found in both flagellar and T3SS operons – for example, in *Luteimonas huabeiensis,* which encodes a flagellar FliO (WP_043689657.1) and a T3SS-associated FliO (WP_024891899.1). In contrast, the closely related *Luteimonas arsenica* harbors only a T3SS-associated FliO (WP_133000184.1) and all canonical flagellar genes are absent. These findings highlight a previously underappreciated diversity of functional contexts for FliO.

### FliO is structurally diverse

Our analysis revealed substantial structural variation among FliO proteins, with lengths ranging from as short as 80 amino acids to over 500 amino acids (Supplementary Data 3). To investigate this diversity, we analyzed representative FliO homologs from diverse phyla using AlphaFold^30^ for structural predictions and applied HHPred to selected longer sequences. These analyses revealed that, despite considerable variation in length and domain architecture, FliO proteins share a conserved structural core. This core structure includes a single transmembrane helix and a small cytoplasmic β-sheet domain (Fig. 3). A few short FliO sequences initially appeared to lack the transmembrane region; however, closer inspection revealed that these cases were likely annotation artifacts, often due to incorrect start codon assignment. For example, *Candidatus Arthromitus* sp. SFB-turkey contains a 60-residue FliO protein in NCBI RefSeq (WP_242861642.1), which lacks a transmembrane segment. However, the corresponding UniProt entry (A0A1B7LMM5) contains an additional 31 N-terminal residues, including a confidently predicted transmembrane helix, with 100% identity in the C-terminal region.

**Fig. 3.**
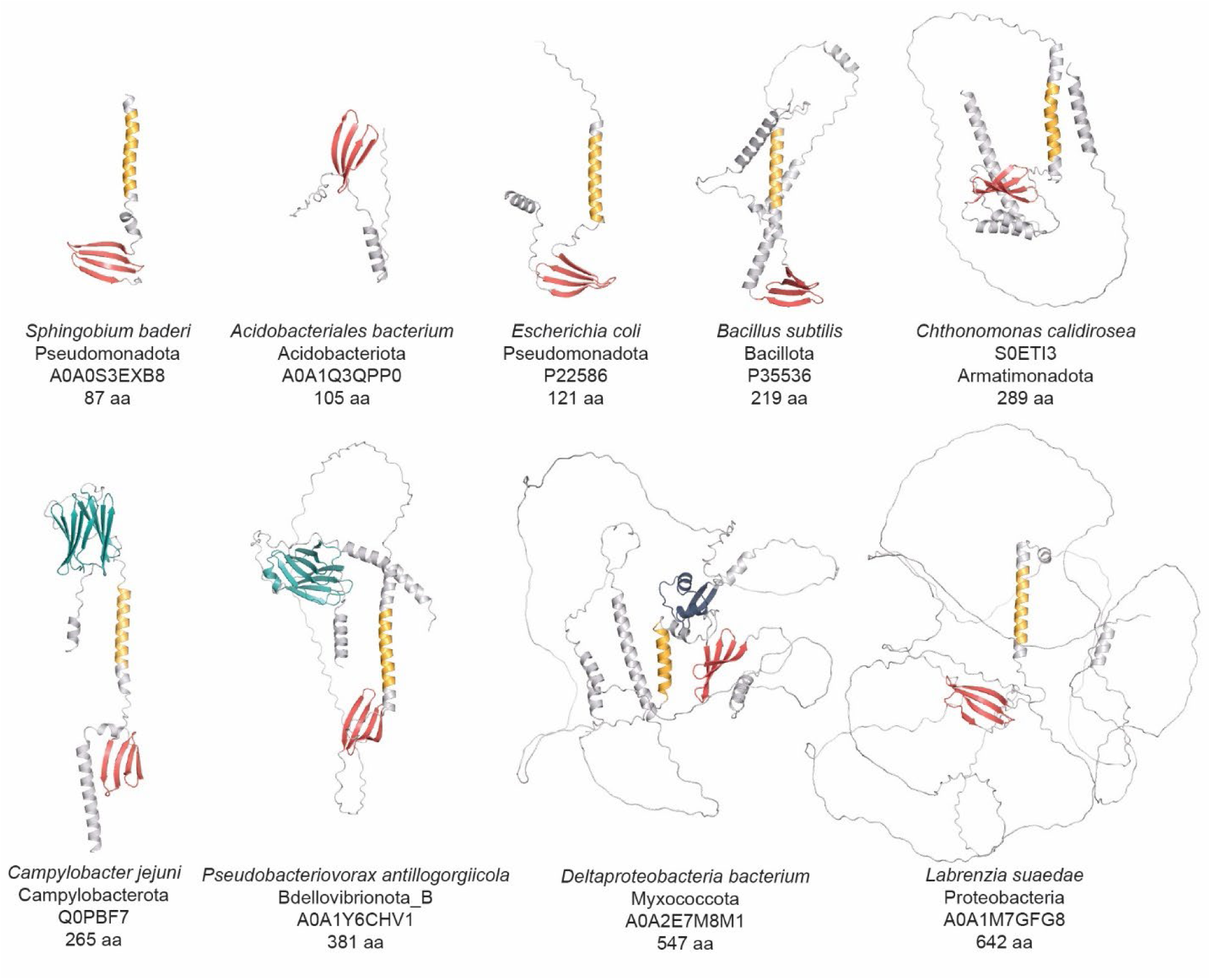
Structural diversity of FliO homologs. Representative structural models of FliO proteins from the AlphaFold Protein Structure Database are shown. Structural elements comprising the conserved core are shown in yellow (a transmembrane helix) and red (an anti-parallel beta-sheet). Variable regions are shown in gray. The AMIN domain is shown in teal, and the LysM domain is shown in blue.

Using Phobius^31^, we confirmed that the β-sheet region is located in the cytoplasm, consistent with its role in the cytoplasmic export apparatus. The majority of FliO proteins analyzed in this study consist solely of this conserved core. However, in some phyla - such as Thermotogota and Acidobacteriota - we identified sequences lacking the transmembrane region (Fig. 3), suggesting either annotation errors or structural divergence. In contrast, the longest FliO proteins, found primarily in alphaproteobacteria, retain the core structure but include additional elements, such as unstructured loops and α-helices (Fig. 3), which may confer phylum-specific functions or interactions.

Intriguingly, FliO proteins from several Myxococcota genomes contain a LysM domain (Fig. 3), a well-characterized peptidoglycan-binding module found in bacterial cell wall enzymes^32^. The presence of LysM suggests a possible link between FliO and cell wall-associated processes in these species.

The most striking example of structural variation was the presence of the AMIN domain in FliO proteins from several phylogenetically diverse bacterial phyla (Fig. 3). The AMIN domain, commonly found in peptidoglycan hydrolases and transport-associated proteins^33^, is essential for localization and substrate interactions. For instance, in *E. coli*, the AMIN domain in AmiC is required for septal ring localization during daughter cell separation^34^. It is also present in PilQ proteins of *Pseudomonas aeruginosa*, *Neisseria meningitidis*, and *Myxococcus xanthus*, where it facilitates transport of type IV pili components across the outer membrane^35,36^.

Recent evidence from *M. xanthus* PilQ suggests that its AMIN domains bind septal and polar peptidoglycan^37^. Consistent with this finding, our Phobius-based analysis predicts that the AMIN domain in FliO is located in the periplasm. We identified N-terminal AMIN domains in FliO proteins from Nitrospinota, Myxococcota, Deferribacterota, Campylobacterota, Bdellovibrionota, and several other phyla (Supplementary Data 2, Fig. 3). Notably, almost all members of the Campylobacterota phylum, including *C. jejuni*, encode FliO proteins containing an AMIN domain.

Taken together, this structural diversity - particularly the acquisition of additional domains such as AMIN - likely contributes to the difficulty of identifying FliO homologs using standard sequence-based approaches. These findings underscore the importance of integrating structure-based analyses to capture the full diversity of the FliO family.

We have chosen the AMIN domain-containing FliO from *Campylobacter jejuni* for experimental analysis to validate computational findings and to understand the role of FliO proteins with this unusual structure.

### FliO and the AMIN domain modestly impact *in vitro* motility and flagellation in *C. jejuni*

Deletion of *fliO* in in *Pseudomonas aeruginosa*, *S. enterica* and *H. pylori* results in severe defects in flagellation and motility^16,17,38^. To assess whether *C. jejuni* FliO plays a similar role, we evaluated the motility and flagellation phenotypes of *C. jejuni* mutants lacking either the entire FliO protein or its AMIN domain. In motility assays using MH soft agar, the Δ*fliO* mutant exhibited a 35% reduction in swimming motility compared to wild-type (WT) *C. jejuni* (Fig. 4A, 4B). Complementation of the Δ*fliO* mutant with either FliO-FLAG or FliO_ΔAMIN_-FLAG variant resulted in a partial, though not statistically significant, restoration of motility.

**Fig. 4.**
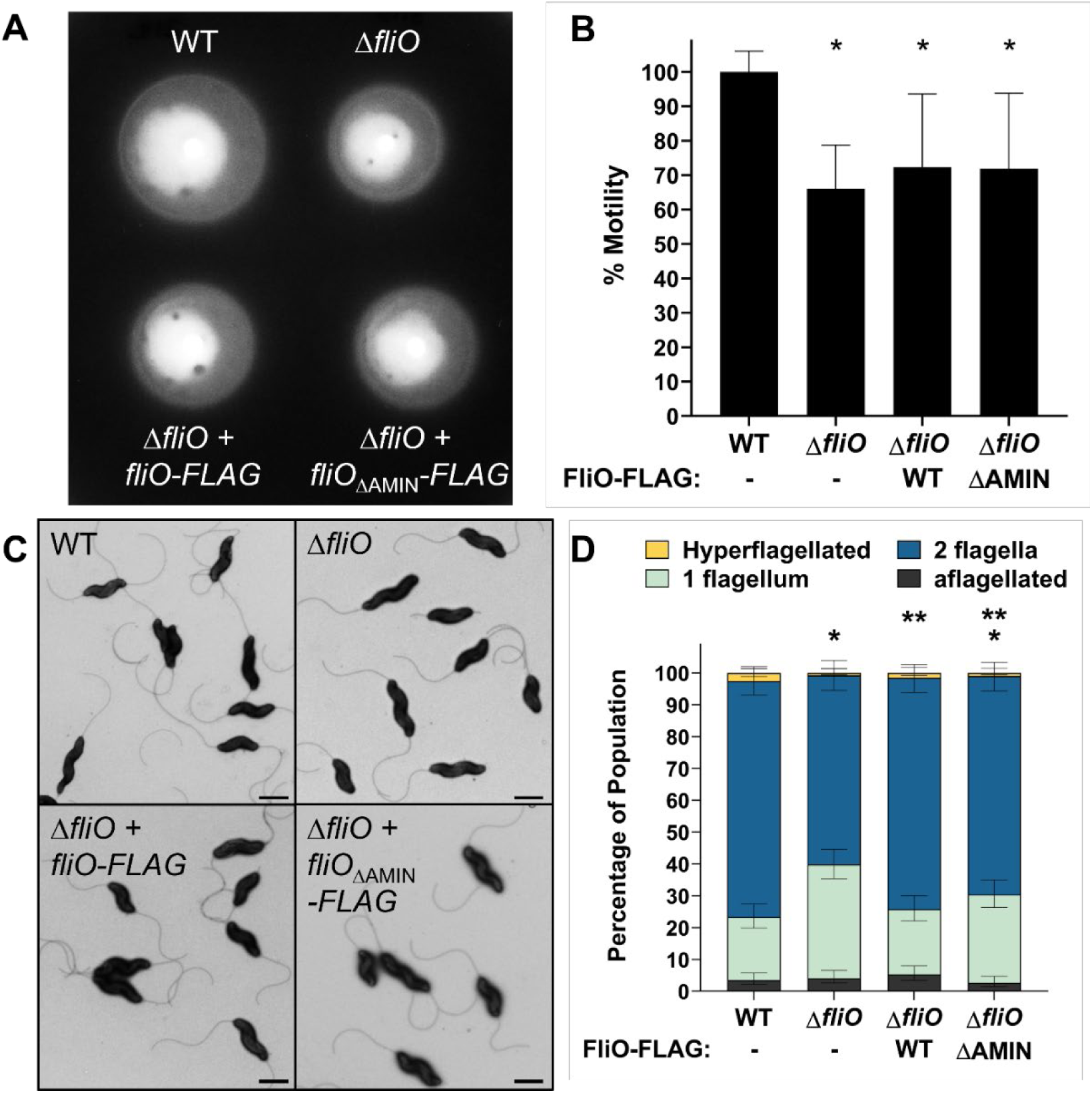
Characterization of the motility and flagellation phenotypes of WT *C. jejuni* and isogenic Δ*fliO* mutants. (A) Representative image of the motility phenotypes of WT *C. jejuni* (top left), Δ*fliO* (top right), Δ*fliO* with *fliO-FLAG* (bottom left), and Δ*fliO* with *fliO*_ΔAMIN_-*FLAG* (bottom right). Strains were diluted to an OD_600_ of 0.8 and then stabbed into MH motility agar containing 0.4% agar. Motility was analyzed after 30 h of incubation at 37°C in microaerobic conditions. (B) Quantitative assessment of the motility phenotypes of each strain. Strains were tested in triplicate. The area of each motile ring was measured, and the average area of each strain was compared to that of WT *C. jejuni,* which was set to 100%. (*, *P* < 0.05 between strains and WT *C. jejuni* as determined by one-way ANOVA with Tukey’s test). (C) Transmission electron micrographs of *C. jejuni* flagellation phenotypes. Strains include WT *C. jejuni* (top left), Δ*fliO* (top right), Δ*fliO* with *fliO-FLAG* (bottom left), and Δ*fliO* with *fliO*_ΔAMIN_-*FLAG* (bottom right). Bar = 1 mm (D) Quantitative assessment of the flagellation phenotype of each strain. Individual bacteria were analyzed by electron microscopy for the number of flagella produced on the poles of each cell. Data are reported as the percentage of the bacterial population that were hyperflagellated (more than one flagellum at either pole), producing two flagella (one flagellum at each pole), producing one flagellum only at one pole, or aflagellated. Three independent populations (n>100) were averaged. Error bars represent standard deviations. Statistical significance was determined by Fisher’s exact test (*, *P* <0.05 between strains and WT *C. jejuni*; **, *P* < 0.05 between strains and Δ*fliO*).

To further investigate the role of FliO in flagellar assembly, we used transmission electron microscopy (TEM) to examine flagellation patterns in WT and mutant *C. jejuni*. Each cell was categorized into one of four phenotypes: (1) hyperflagellated (>1 flagellum at a pole), (2) bipolar flagellation (1 flagellum per pole), (3) unipolar flagellation (1 flagellum at a single pole), or (4) aflagellated (no flagella). Over 90% of cells in all strains produced at least one flagellum (Fig. 4C, 4D). However, the Δ*fliO* mutant showed a notable shift in flagellation pattern, with a decreased proportion of cells producing flagella at both poles (∼59% in Δ*fliO* vs. ∼74% in WT) and an increased proportion of unipolar flagellated cells (∼36% in Δ*fliO* vs. ∼20% in WT) (Fig. 4C, 4D). This suggests that FliO promotes efficient bipolar flagellation, although it is not essential for flagellum formation. Complementation with FliO-FLAG restored the WT flagellation pattern, whereas FliO_ΔAMIN_-FLAG only partially restored it, with ∼69% of cells exhibiting bipolar flagellation and ∼27% displaying unipolar flagella (Fig. 4C, 4D). These results imply that the AMIN domain contributes to - but is not solely responsible for - FliO’s function in flagellar assembly.

### *C. jejuni* FliO influences the stability of FlhB but not other fT3SS proteins

In *Salmonella*, FliO functions as a chaperone that stabilizes FliP and facilitates the formation of the FliP–FliR complex^16^. Consequently, *fliO* deletion in *Salmonella* leads to increased instability of FliP^15,16^. To investigate whether *C. jejuni* FliO plays a similar role, we analyzed FliP protein levels in total membrane fractions from WT and Δ*fliO* strains. Due to the lack of reliable anti-FliP antisera, we constructed a chromosomal *fliP*-FLAG allele, inserting a FLAG epitope in-frame immediately downstream of the predicted N-terminal signal peptide (between codons 22 and 23). Production of FliP-FLAG by *C. jejuni* allowed for a significant level of motility comared to *C. jejuni* Δ*fliP*, but motility of this strain was modestly lower than WT (Fig. S2). Immunoblot analysis revealed that FliP-FLAG levels were comparable between WT and *ΔfliO* (Fig. 5A). These findings were corroborated by mass spectrometry of membrane fractions (Table. S3). Thus, FliO is not required for steady-state levels of FliP in *C. jejuni*.

**Fig. 5.**
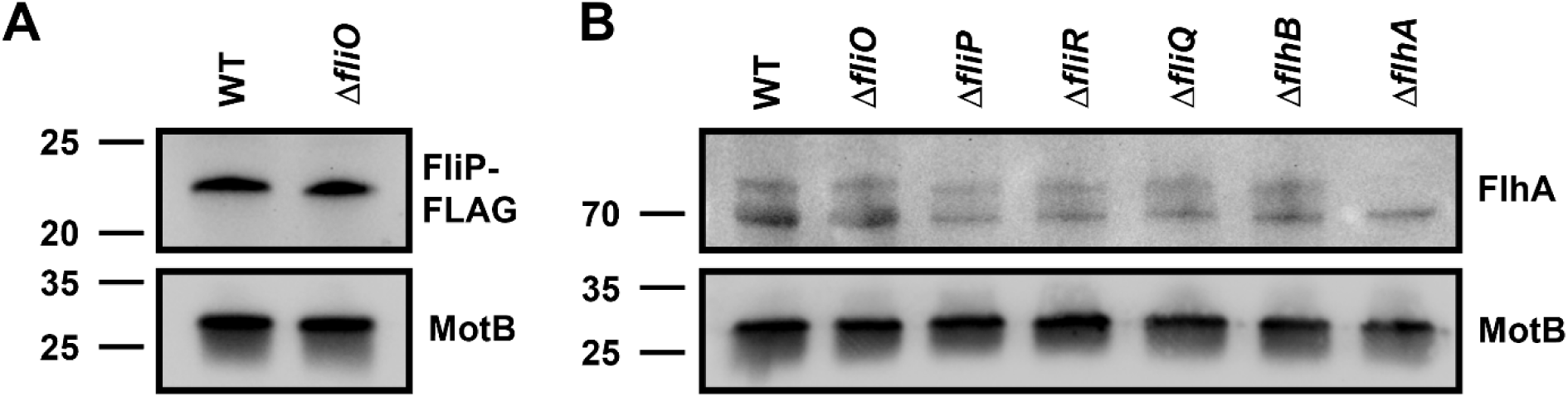
FliP and FlhA levels in WT *C. jejuni* and isogenic mutants. (A) Representative immunoblot analysis of FliP-FLAG production in the total membrane fraction of WT *C. jejuni* and the Δ*fliO* mutant. Equal levels of membrane proteins (5 mg) from each strain were examined. a-FLAG antibody was used to detect FliP-FLAG (B) Representative immunoblot analysis of FlhA in the total membrane fraction of WT *C. jejuni* and fT3SS mutants. Equal levels of membrane proteins (10 mg) from each strain were examined. Custom a-FlhA antiserum was used to detect FlhA. Detection of MotB served as a control to ensure equal loading of total membrane proteins across strains in both A and B. Specific antiserum was used to detect *C. jejuni* MotB.

In *H. pylori*, deletion of *fliO* resulted in a 2- to 3-fold reduction in FlhA levels, rather than FliP^17^. We therefore assessed FlhA protein levels in membrane fractions from WT *C. jejuni*, Δ*fliO*, and additional fT3SS mutants (Δ*fliQ*, Δ*flhB*), using FlhA-specific antisera previously developed in our lab^39^. Unlike in *H. pylori*, FlhA levels remained unchanged in Δ*fliO*, Δ*fliQ*, and Δ*flhB* strains (Fig. 5B), consistent with results from mass spectrometry (Fig. S3). In contrast, FlhA levels were reduced by ∼25% in Δ*fliP* and Δ*fliR* mutants (Fig. 5B). Overall, our analysis does not implicate FliO involvement in maintaining significant stability of FlhA, unlike in *H. pylori*. These findings also imply that FliO is not required for FliR stability.

Next, we examined whether *C. jejuni* FliO influences the stability of other fT3SS components. Although we were unable to detect FliQ or FliR—due to lack of antisera and unsuccessful FLAG-tagging—we successfully measured FlhB levels using specific antisera developed previously^40^. In Δ*fliO* strains, FlhB levels were reduced by approximately 40% compared to WT (Fig. 6A, 6B), a result confirmed by mass spectrometry (Table. S3). Notably, complementation with FliO-FLAG restored FlhB levels, whereas FliO_ΔAMIN_-FLAG failed to do so, indicating that the AMIN domain is required for FlhB stabilization.

**Fig. 6.**
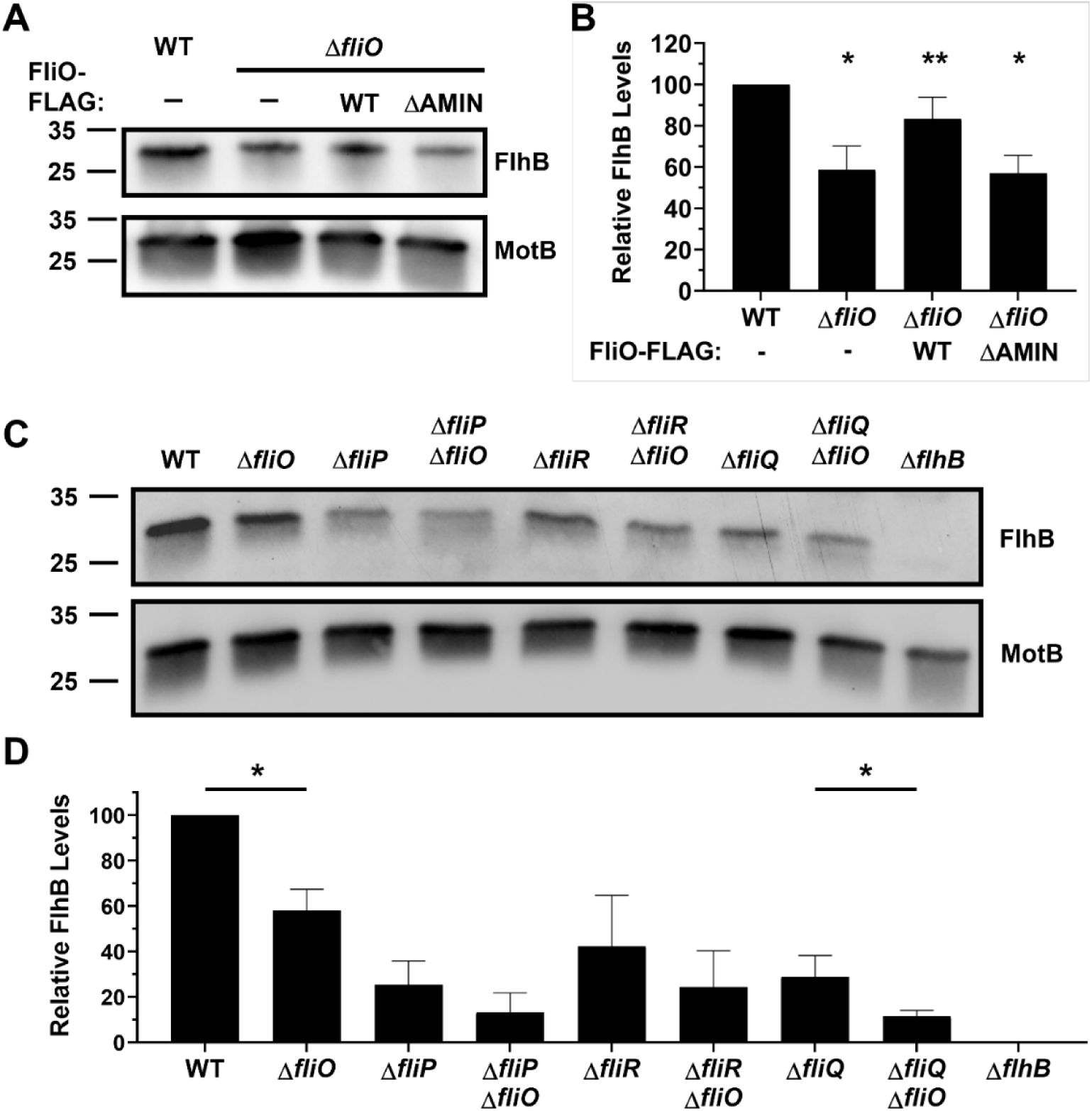
FlhB levels in WT *C. jejuni* and isogenic mutants. (A) Immunoblot analysis of FlhB in the total membrane fraction of WT *C. jejuni,* Δ*fliO*, Δ*fliO* with *fliO-FLAG*, and Δ*fliO* with *fliO*_ΔAMIN_-*FLAG*. Equal amounts of membrane proteins (5 mg) from each strain were examined. Specific antisera were used to detect FlhB and MotB, which served as a control to ensure equal loading of proteins. (B) Quantitative assessment of FlhB levels by densitometry. Immunoblots from three individual preparations of membranes from strains were examined (with one immunoblot represented in A) and normalized to MotB levels. Statistical analysis was determined by a two-tailed unpaired *t*-test (*, *P* < 0.05 between strains and WT *C. jejuni*; **, *P* < 0.05 between strains and Δ*fliO*). (C) Immunoblot analysis of FlhB in the total membrane fraction of WT *C. jejuni* and mutants lacking one or more components of the fT3SS. Equal levels of total membrane proteins (5 mg) were examined. Specific antisera were used to detect FlhB and MotB, which served as a control to ensure equal loading of proteins (D) Quantitative assessment of FlhB levels by densitometry. Immunoblots from three individual preparations of membranes from strains were examined (with one immunoblot represented in C) and normalized to MotB levels. Statistical analysis was determined by a two-tailed unpaired *t*-test (*, *P* < 0.05 between pairs of strains with and without *fliO*).

To assess whether the effect of FliO on FlhB is independent of other fT3SS proteins, we compared FlhB levels in Δ*fliP*, Δ*fliR*, and Δ*fliQ* mutants—with and without *fliO*. In these mutants, FlhB levels were already reduced to ∼25–40% of WT, consistent with the idea that secretion pore assembly is necessary for FlhB stability (Fig. 6C, 6D). Additional deletion of *fliO* in these mutants further reduced FlhB levels, although the difference reached statistical significance only in the Δ*fliQ* Δ*fliO* strain (Fig. 6C, 6D).

Collectively, these data indicate that *C. jejuni* FliO may act upon FlhB independent of FliP, FliR, and FliQ to maintain levels of FlhB. Furthermore, the requirement of the AMIN domain for this function suggests a structural role in stabilizing FlhB within the membrane.

### *C. jejuni* FliO and the FliO AMIN domain are necessary for WT levels of commensal colonization of chicks

*C. jejuni* is a commensal intestinal bacterium in many animal hosts - particularly birds - and a leading cause of bacterial gastroenteritis in humans^41–44^. Previous studies have demonstrated that flagellar motility is essential for *C. jejuni* colonization of both avian and human intestinal tracts and is critical for epithelial cell interaction and invasion^42,45,46^.

To assess the role of FliO and its AMIN domain *in vivo*, we analyzed commensal colonization of the chick cecum using WT *C. jejuni*, a Δ*fliO* mutant, and Δ*fliO* mutants complemented with FliO-FLAG or FliO_ΔAMIN_-FLAG. The complementation constructs were integrated into a non-native chromosomal locus, driven by the native *fliNO* promoter. Although this promoter is relatively weak and precluded detection of FliO-FLAG and FliO_ΔAMIN_-FLAG in whole-cell lysates, we confirmed comparable expression of both proteins by immunoprecipitation with anti-FLAG resin followed by immunoblotting (Supplementary Fig. 1).

Day-old chicks were orally inoculated with ∼100 CFU of each strain. Seven days post-infection, bacterial loads in the cecal contents were quantified. As previously reported^47,48^, WT *C. jejuni* colonized the ceca at a level of ∼1.2 × 10⁹ CFU/g (Fig. 7). In contrast, the *ΔfliO* mutant exhibited an 8.2-fold reduction in colonization. Complementation with fliO-FLAG fully restored colonization to WT levels, confirming that the colonization defect was due to loss of FliO. However, complementation with fliO_ΔAMIN_-FLAG failed to rescue the phenotype, and colonization levels remained comparable to the Δ*fliO* mutant.

**Fig. 7.**
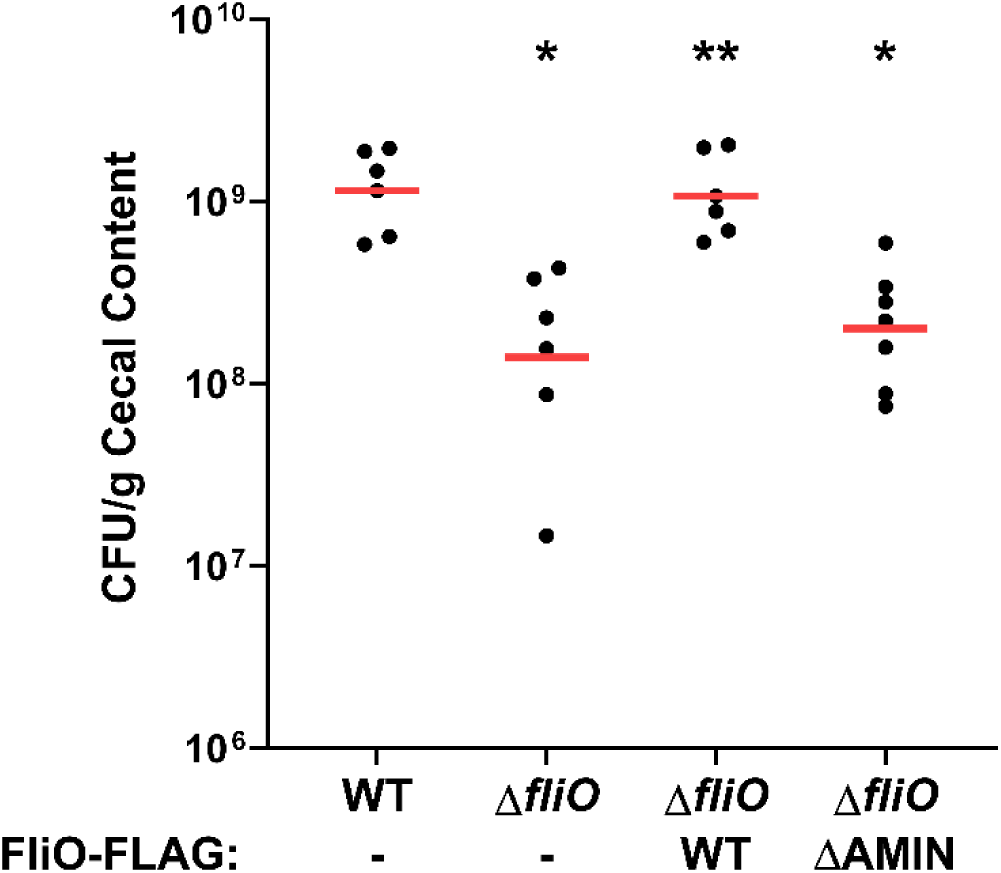
Commensal colonization of WT *C. jejuni* and isogenic Δ*fliO* mutants in avian hosts. One-day old chicks were orally gavaged with ∼100 colony forming units (CFU) of *C. jejuni* strains. At day 7, chicks were sacrificed and the CFU per gram of cecal content were determined. Each circle represents the CFU counts per g of cecal content from an individual chick. Red bars represent the geometric mean for each group. Statistical analysis was performed using the Mann-Whitney *U* test (*, *P* < 0.05 between strains and WT *C. jejuni*; **, *P* < 0.05 between strains and Δ*fliO)*.

Together, these results demonstrate that FliO is essential for efficient colonization of the chick intestinal tract, and that the AMIN domain plays a critical role in mediating this function. The data further support a role for FliO in *C. jejuni* colonization that is at least partially distinct from that of FliO homologs in other bacterial species.

## Discussion

This study provides the most comprehensive analysis to date of phyletic distribution, structural and functional diversity, and evolution of the flagellar protein FliO, a core component of the fT3SS.

Earlier studies suggested a sporadic distribution of FliO across bacterial species^6,16^, and consequently its evolutionary history remained unknown, and it is not considered as a core component of the bacterial flagellar system. Using a highly sensitive, curated computational pipeline, we show that FliO is present in **∼**95% of genomes encoding core flagellar components FliP, FliQ, and FliR, indicating that FliO likely originated in the last common ancestor of flagellated bacteria. Its absence is best explained by secondary losses, observed in limited lineages within only two phyla, such as *Arcobacteraceae* (Campylobacterota) and certain Pseudomonadota (e.g., *Rhodobacterales*, *Caulobacterales*). These rare absences strongly suggest lineage-specific adaptations and imply that FliO was present in the fT3SS of the last common ancestor of bacteria, which is thought to have been flagellated^26^.

Unexpectedly, we also identified FliO homologs in non-flagellar T3SS, particularly in Myxococcota, where FliO is associated with predation-related systems^29^, as well as in Pseudomonadota including *Xanthomonadales* and *Caulobacterales*. Moreover, FliO homologs were found in non-flagellated organisms such as *Chlamydiia* (Verrucomicrobiota A), where they are encoded in conserved operons of unknown function. These observations suggest that FliO may have been repurposed for secretion systems beyond motility, supporting the growing recognition of modular and transferable architectures in bacterial secretion systems^28^.

A major outcome of our study is the revelation of structural diversity among FliO proteins. While most FliO homologs consist solely of a conserved core - comprising a single transmembrane segment and a cytoplasmic β-sheet - a subset contains additional structural features, including entire domains such as LysM and AMIN. The LysM domain is found in a wide variety of extracellular proteins and is known for a general peptidoglycan binding function^32,49^. The AMIN domain mediates localization of diverse periplasmic protein complexes^33^. In *E. coli* cell division amidase AmiC, the N-terminal AMIN domain binds to peptidoglycan and is involved in its localization at the division site^34^. This domain is also found in PilQ secretin, a component of the conserved Type IVa pili (T4aP) machinery, where it binds peptidoglycan and ensures proper localization and stabilization of the PilQ secretin complex^37^. Recent studies revealed novel structural elements in the flagellar motor of *H. pylori* and *C. jejuni* including the AMIN domain-containing motor scaffold protein PflA^50,51^. The presence of AMIN domain in FliO from *Campylobacterota* and representative of several other phyla strongly suggests an evolutionarily acquired role in envelope anchoring.

Experimental analysis in *C. jejuni* revealed that FliO contributes to proper flagellation and motility, but its deletion resulted in milder phenotypes compared to *S. enterica* and *H. pylori*, where *fliO* deletion leads to severe aflagellation^17,52^. In *C. jejuni*, the Δ*fliO* mutant retained flagella at one or both poles in over 90% of cells, though bipolar flagellation and swimming motility were reduced. The AMIN domain was necessary to fully restore the wild-type phenotype, implicating it in fine-tuning polar flagella formation or positioning.

Importantly, we found little evidence to support that FliO is required for the stability of FliP or FlhA, contrasting with prior work in *Salmonella* and *H. pylori*^15–17^. However, FliO significantly influences the membrane stability of FlhB, a core export gate protein, supporting a chaperone-like role specific to FlhB in *C. jejuni*. This stabilizing effect is dependent on the AMIN domain, highlighting the functional importance of its acquisition.

FliO was also required for efficient colonization of the chick cecum, a model of *C. jejuni* commensalism. Deletion of *fliO* led to an 8-fold reduction in colonization, which was fully rescued by complementation with WT fliO but not with a version lacking the AMIN domain. These results establish a link between flagellar assembly, domain-level modularity in flagellar accessory proteins, and host colonization.

Together, our findings establish FliO as a core, yet evolutionarily dynamic, component of the bacterial flagellum - and, in some cases, of related secretion systems. The expansion of domain architecture (e.g., LysM, AMIN) reflects broader principles of modularity and innovation in bacterial envelope biology. Contextual repurposing of FliO supports the notion that flagellar motor assembly is a dynamic process in which accessory proteins may interact transiently with other components of this complex nanomachine^53^. While prior knowledge on FliO was based on studies in only a few species and limited phylogenomic analyses, our data suggest that FliO functions may vary across species, shaped by lineage-specific regulatory and structural constraints. Challenges with FliO detection within genomic datasets due to substantial sequence variability indicate that some other flagellar components may in fact be integral parts of the core machinery but are currently overlooked. Finally, this study and other recent findings on loss and acquisition of flagellar genes^54,55^ strongly suggest that evolution of bacterial flagellum is more complex than previously thought.

## Materials and Methods

### Bioinformatics tools and databases

BLAST and PSI-BLAST searches^18^ against selected genomes were performed with default parameters. Taxonomy tree and information for phyletic distribution were retrieved from the Genome Taxonomy Database (GTDB), v. 95.0^22^. Protein sequences of a representative set of bacterial genomes were downloaded from the NCBI RefSeq, NCBI nr^56^ and GTDB databases. The retrieved dataset was scanned with the KEGG^25^ profile HMMs: K02418, K02419, K02420, K02421 for FliO, FliP, FliQ, and FliR, respectively, and Pfam^21^ PF04347 for FliO with the E-value thresholds set either to default or to 500 (a relaxed threshold). Domains were identified using the TREND resource^57,58^ with the Pfam profile HMMs (v. 35.0). Gene neighborhoods were identified using TREND and MiST4 database^59^. The presence of the FliO domain in selected sequences was verified using HHpred^24^. Predicted structures of FliO proteins were retrieved from the AlphaFold structural database^30^ and visualized in PyMOL 3.0. Transmembrane regions, signal peptides, and cellular localization were determined using Phobius^31^.

### Bacterial Strains and Growth Conditions

All bacterial strains created and used in this study are described in Supplementary Table 1. All plasmids constructed or involved in creation of strains used in this study are described in Supplementary Table 2.

*E. coli* DH5α and DH5α/pRK212.1 were used for plasmid construction and conjugation of plasmids into *C. jejuni*, respectively ^60^. All *E. coli* strains were grown in Luria-Bertani (LB) broth or on LB agar with appropriate antibiotics (100 mg/mL ampicillin, 15 µg/mL chloramphenicol, 50 µg/mL kanamycin, or 12.5 µg/mL tetracycline) at 37 °C. *E. coli* strains were stored at −80°C in an 80% LB broth and 20% glycerol solution.

For all experiments, *C. jejuni* strains were grown from freezer stocks for 48 h and then restreaked and grown for 16 h at 37 °C in microaerobic conditions (85% N_2_, 10% CO_2_, and 5% O_2_) on Mueller-Hinton (MH) agar supplemented with appropriate antibiotics (10 mg/mL trimethoprim, 30 μg/mL cefoperazone, 10 mg/mL chloramphenicol, 50 mg/mL kanamycin, or 0.1, 0.5, 1.0, or 2.0 mg/mL streptomycin). *C. jejuni* strains were stored at −80 °C in an 85% MH broth and 15% glycerol solution.

### Generation of *C. jejuni* Mutants

All *C. jejuni* mutants were generated by electroporation of plasmid DNA following previously described methods^61,62^. All plasmids used to create *C. jejuni* mutants were constructed by ligation of DNA fragments into plasmids by T4 DNA ligase or Gibson Assembly Master mix (New England BioLabs).

Primers were designed to amplify the *fliO* locus from 0.7 kb upstream and downstream of *fliO* from the genome of *C. jejuni* 81-176. The DNA fragment was cloned into the BamHI site of pUC19 by Gibson assembly to create pDRH2455. Primers were designed to amplify the *fliO* locus from pDRH2455 to contain a T410G mutation which created an StuI site within *fliO*. The DNA fragments were cloned into the BamHI site of pUC19 by Gibson assembly to create pDRH2559. The SmaI-digested *cat-rpsL* cassette from pDRH265 was inserted into the StuI site of *fliO* in pDRH2559 to create pDRH2568 ^63^. pDRH2568 was introduced into 81-176 *rpsL*^Sm^ and transformants were recovered on MH agar with chloramphenicol. Transformants were verified by colony PCR to result in DRH8073 (81-176 *rpsL*^Sm^ *fliO::cat-rpsL*). Primers were designed to amplify the *fliO* locus from pDRH2455 and generate in-frame deletion of *fliO* that fused the start codon of *fliO* to codon 264 to create pDRH2547. pDRH2547 was transformed into DRH8073 and transformants were recovered on MH agar with streptomycin. Transformants were screened for chloramphenicol-sensitivity and then verified by colony PCR to result in DRH8113 (81-176 *rpsL*^Sm^ D*fliO*).

For complementation of *C. jejuni* Δ*fliO*, constructs encoding *fliO-FLAG* or *fliO*_ΔAMIN_-*FLAG* under expression by the native *fliNO* promoter were cloned into the *C. jejuni rrsC*-*rrlC* locus. First, a 2.6 kb DNA fragment containing a portion of the coding sequence of *rrsC* through a portion of *rrlC* was cloned into the EcoRI site of pBR322 to create pDRH7574. Primers were designed to amplify the *kan* cassette from pILL600 such that the XbaI restriction site at the 3’ end remained intact ^64^. The *kan* cassette was cloned into the XbaI site of the *rrsC-rrlC* locus in pDRH7574 by Gibson assembly to create pDRH7746. Primers were designed to amplify DNA fragments containing the *fliNO* promoter from 120 bp up to the *fliN* start codon fused to the *fliO* coding sequence with a C-terminal FLAG tag. These DNA fragments were cloned into the XbaI site of pDRH7746 by Gibson assembly to create pALD1047. Additional primers were designed to make a similar construct that created a fusion of codons 20 and 119 of *fliO* to delete the predicted AMIN domain to result in pALD1059. pALD1047 and pALD1059 were transformed into 81-176 *rpsL*^Sm^ Δ*fliO* and transformants were recovered on MH agar with kanamycin. Mutants were verified by colony PCR to result in ALD1050 (81-176 *rpsL*^Sm^ Δ*fliO rrsC::kan-P_fliNO_-fliO-FLAG*) and ALD1081 (81-176 *rpsL*^Sm^ Δ*fliO rrsC::kan-P_fliNO_-fliO*_DAMIN_*-FLAG*).

Primers were designed to amplify the *fliP* locus from 0.7 kb upstream and downstream of *fliP* from the genome of *C. jejuni* 81-176 with a FLAG tag encoded after the predicted N-terminal signal sequence of FliP between codons 22 and 23 of the gene. The DNA fragments were cloned into the EcoRI site of pUC19 by Gibson assembly to create pALD1062 (pUC19::*fliP_P22-FLAG-T23_*). To replace *fliP* on the chromosome of a Δ*fliO* mutant, DRH8113 (81-176 *rpsL*^Sm^ Δ*fliO*) was first transformed with pDRH643^63^. Transformants were recovered on MH agar with chloramphenicol and verified by colony PCR to obtain ALD963 (81-176 *rpsL*^Sm^ Δ*fliO fliP::cat-rpsL*). pALD1062 was transformed into DRH706 (81-176 *rpsL*^Sm^ *fliP::cat-rpsL*) or ALD963 (81-176 *rpsL*^Sm^ Δ*fliO fliP::cat-rpsL*) and transformants were recovered on MH agar containing streptomycin ^63^. Chloramphenicol-sensitive transformants were verified by colony PCR for replacement of WT *fliP* with *fliP_P22-FLAG-T23_* to obtain ALD1130 (81-176 *rpsL*^Sm^ *fliP_P22-FLAG-T23_*) and ALD1106 (81-176 *rpsL*^Sm^ Δ*fliO fliP_P22-FLAG-T23_*).

To generate *fliR* and *fliQ* mutants in *C. jejuni* Δ*fliO*, DRH8113 was transformed with pDRH645 and pSMS469 and transformants were recovered on MH agar with kanamycin or chloramphenicol, respectively^63,65^. Transformants were verified by colony PCR to result in ALD962 (81-176 *rpsL*^Sm^ Δ*fliO fliR::kan-rpsL*) and ALD965 (81-176 *rpsL*^Sm^ Δ*fliO fliQ::cat-rpsL*).

### Chick Colonization Assays

All chicken colonization experiments were approved by IACUC at the University of Texas Southwestern Medical Center. Assays to determine the ability of WT *C. jejuni* 81-176 *rpsL*^Sm^ and isogenic mutants to colonize the chick ceca following oral gavage were performed as previously described^45^. Briefly, fertilized chicken eggs (Charles River Avian Vaccine Services) were obtained and incubated in the Sportsman II model 1502 incubator (Georgia Quail Farms Manufacturing Company) for 21 d at 37.8 °C with appropriate humidity and rotation. On the day of hatch, chicks were orally gavaged with 100 μL of PBS containing ∼10^2^ CFU of WT *C. jejuni* 81-176 *rpsL*^Sm^ and isogenic mutants. Dilutions of the inoculum were plated on MH agar supplemented with trimethoprim and cefoperazone to determine the CFUs in each inoculum. At 7 d post-infection, chicks were sacrificed and the cecal contents were recovered, weighed, and suspended in PBS to 0.1 g cecal content/mL. Serial dilutions were plated on MH agar supplemented with trimethoprim and cefoperazone. Bacteria were grown for 72 h at 37 °C in microaerobic conditions and then colonies were counted to determine the CFU per gram of cecal content for each chick. Recovered colonies were analyzed by colony PCR to confirm recovered bacteria matched the strain with which each chick was infected. Statistical analyses were performed in GraphPad Prism using a Mann-Whitney *U* test, with statistically significant differences indicating *P* < 0.05.

### Immunoprecipitation and detection of *C. jejuni* FliO-FLAG and FliO_DAMIN_-FLAG

Immunoprecipitation of FliO-FLAG or FliO_ΔAMIN_-FLAG from *C. jejuni* strains was performed as previously described^66,67^. *C. jejuni* strains were grown overnight for 16 h at 37 °C in microaerobic conditions on MH agar plates supplemented with 50 mg/mL kanamycin. Strains were resuspended in MH broth and diluted to OD_600_ of 0.8. Twenty milliliters of each culture were pelleted, washed in PBS, and then fixed with formaldehyde. After incubating for 30 min at 37 °C, glycine was added to a final concentration of 160 mM and incubated for 10 min at 25 °C. After one wash in PBS, cells were lysed by osmolysis. Lysates were solubilized by the addition of a solution containing 50 mM Tris, pH 8.0, 10 mM MgCl_2_, and 2% Triton X-100. After incubation, the lysate was pelleted for 20 min at 10,000 rpm. The supernatant was mixed with anti-FLAG affinity resin (Sigma-Aldrich) and incubated overnight at 4 °C. The resin was pelleted for 10 min at 13,000 rpm at 4 °C, washed 3 times with radioimmunoprecipitation assay (RIPA) buffer (50 mM Tris, pH 8.0, 150 mM NaCl, 0.1% SDS, 0.5% sodium deoxycholate, 1% Triton X-100) and resuspended in sodium dodecyl sulfate-polyacrylamide gel electrophoresis (SDS-PAGE) loading buffer containing 0.5% β-mercaptoethanol. FliO-FLAG and FliO_ΔAMIN_-FLAG were detected by immunoblot analysis as described below.

### Motility Assays

*C. jejuni* strains were grown overnight for 16 h at 37 °C in microaerobic conditions on MH agar with appropriate antibiotics. Strains were resuspended and then diluted to OD_600_ of 0.8 and then stabbed into MH agar containing 0.4% agar and trimethoprim. Motility was analyzed after 30 h of incubation at 37 °C under microaerobic conditions. Motility rings were imaged with the ChemiDoc Imaging System (Bio-Rad) and analyzed with the Image Lab (Bio-Rad) software. Areas of motility ring areas from three independent cultures were averaged and normalized to that of WT *C. jejuni*. Statistical analyses were performed in GraphPad Prism using a two-tailed Student’s *t* test, with statistically significant differences indicating *P* < 0.05.

### Transmission Electron Microscopy

Flagellation phenotypes for WT *C. jejuni* 81-176 *rpsL*^Sm^ and isogenic mutants were analyzed by transmission electron microscopy (TEM). *C. jejuni* strains were grown overnight for 16 h at 37 °C in microaerobic conditions on MH agar with appropriate antibiotics. Strains were diluted to an OD_600_ of 0.8. One mL of each culture was pelleted for 3 min at 13,000 rpm, resuspended in 2.5% glutaraldehyde in 0.1 M cacodylate, and then incubated on ice for 1 h. Bacterial samples were applied to negatively glow-discharged copper-coated formvar grids. The samples were stained with 2% uranyl acetate and visualized with the JEOL 1400 Plus transmission electron microscope. Electron micrographs were analyzed using Fiji (ImageJ2). For each culture, over 100 individual cells were analyzed to determine if a cell was aflagellated (0 flagella per cell), produced one flagellum (only 1 flagellum at one of the poles), produced two flagella (a single flagellum at both poles), or was hyperflagellated (produced more than one flagellum at one of the poles). Results from three independent cultures were averaged to determine the percentage of the population that produced each flagellation phenotype. Statistical analyses were performed in GraphPad Prism using a Fisher’s exact test, with statistically significant differences indicating *P* < 0.05.

### Fractionation of *C. jejuni* cells for protein analysis

Whole-cell lysates (WCL) and total membrane fractions were isolated to assess levels of flagellar proteins in WT *C. jejuni* 81-176 *rpsL*^Sm^ and isogenic mutants, as previously described^40^. Briefly, *C. jejuni* strains were grown overnight for 16 h at 37 °C in microaerobic conditions on MH agar plates supplemented with appropriate antibiotics. Strains were resuspended and diluted to OD_600_ of 0.8 in MH broth. To isolate WCLs, 1 mL of each culture was pelleted for 3 min at 13,000 rpm, washed once in PBS, and resuspended in SDS-PAGE loading buffer containing 0.5% β-mercaptoethanol. To isolate total membrane fractions, 5 mL of each culture was pelleted for 10 min at 6,000 rpm, washed once in 1 mL of 10 mM HEPES (pH 7.4), and resuspended in 1 mL of 10 mM HEPES. Cells were lysed via sonication using the Branson Sonifier 450 set at an amplitude of 4.5 with a constant duty cycle. Total membrane was pelleted for 30 min at 13,000 rpm. The pellets were washed three times in 10 mM HEPES and resuspended in SDS-PAGE loading buffer containing 0.5% β-mercaptoethanol.

### Immunoblot analyses

Protein samples were boiled for 5 min and loaded on appropriate percentage SDS-PAGE gel and then transferred to a PVFD membrane using the Trans-Blot Turbo transfer system (Bio-Rad). Blots were blocked with PBS containing 5% powdered milk and 0.1% Tween-20 overnight at 4 °C. Primary antisera were used at the following concentrations: FlhA GP153, 1:100^39^; FlhB Rab476, 1:500^40^; MotB GP140, 1:500^68^; FLAG, 1:500 (Sigma). Blots were incubated with primary antisera for 1 h up to overnight. Secondary HRP-conjugated antibodies were used at a dilution of 1:10,000 and incubated for 1 hr at RT. Blots were developed using the Western Lightning Plus ECL Kit (Perkin-Elmer) and then imaged with the ChemiDoc Imaging System (Bio-Rad). Band densitometry was performed with Image Lab software (Bio-Rad). The statistical significance of the difference in relative band densitometry between samples was determined with a two-tailed Student’s *t* test, with statistically significant differences indicating *P* < 0.05.

### Mass spectrometry analysis of membrane protein composition

Total membrane fractions were isolated to assess levels of flagellar proteins in WT *C. jejuni* and Δ*fliO* as described above. Equal amount of total membrane proteins from each strain were run on an SDS-PAGE gel for 5-10 min and stained with Coomassie Brilliant Blue. Stained band mass representing total membrane protein was excised from gels and submitted to the UTSW Proteomics Core for identification and quantification of *C. jejuni* 81-176 proteins. MS data were acquired using an Orbitrap Fusion Lumos mass spectrometer (Thermo Fisher) coupled to an Ultimate 3000 RSLC-Nano liquid chromatography systems (Thermo Fisher). Raw MS data files were analyzed using Proteome Discoverer v3.0 SP1 (Thermo), with peptide identification performed using Sequest HT searching against the *Campylobacter jejuni* 81-176 protein database from UniProt. The false-discovery rate (FDR) cutoff was 1% for all peptides. Peptide peak intensities were summed for all peptides matched to a protein for protein abundance values. Protein abundance values were normalized such that the sums of the protein abundances for each sample were equal.

## Data availability

All data supporting the findings of this study are available within the paper and its Supplementary Information.

## Acknowledgements

This work was supported, in part, by the National Institutes of Health grants R35GM131760 (to I.B.Z), R01AI065539 (to D.R.H.), and F31AI186490 (to A.L.D) and by the Deutsche Forschungsgemeinschaft (DFG) research grant no. ER 778/2-1, the European Research Council under the European Union’s Horizon 2020 research and innovation program (grant agreement no. 864971) and a Max Planck Fellowship of the Max Planck Society (to M.E.).

## Author contributions

Conceptualization: I.B.Z, M.E., and D.R.H.

Investigation: E.P.A., A.L.D., and I.B.Z.

Visualization: E.P.A. and A.L.D.

Supervision: I.B.Z. and D.R.H.

Writing - original draft: E.P.A. and A.L.D.

Writing – review and editing: I.B.Z., M.E., and D.R.H.

## Competing interests

The authors declare that they have no competing interests.

## Notes

### Competing Interest Statement

The authors have declared no competing interest.

