## Supplementary Figures and Tables for "FliO is an evolutionarily conserved yet diversified core component of the bacterial flagellar type III secretion system"

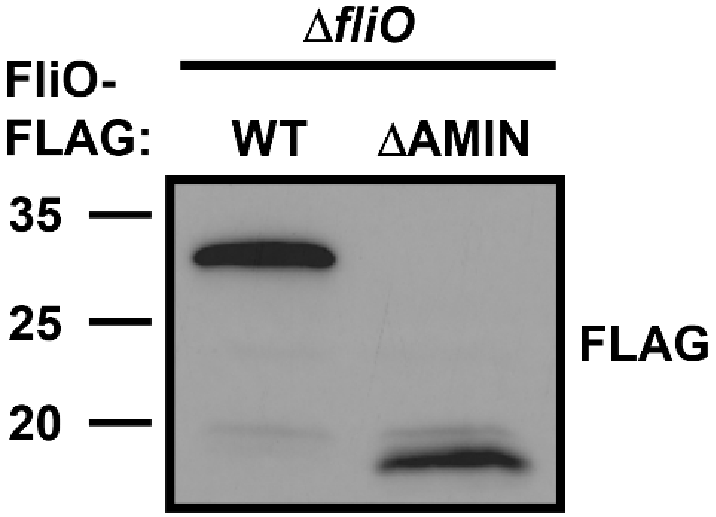

**Supplementary Fig. 1: Immunoblot analysis of immunoprecipitated WT FliO-FLAG and FliO$_{\Delta AMIN}$-FLAG.** *C. jejuni* Δ*fliO* was complemented in the *rrsC* locus with the native *fliNO* promoter expressing *fliO-FLAG* or *fliO*$_{\Delta AMIN}$-*FLAG*. Strains were grown and then diluted to $OD_{600}$ of 0.8. Proteins were immunoprecipitated with a-FLAG resin and, after washing, resuspended in equal volume of 1X SDS-PAGE loading buffer. Equal volumes of samples were examined. a-FLAG antibody was used to detect FliO-FLAG and FliO$_{\Delta AMIN}$-FLAG.

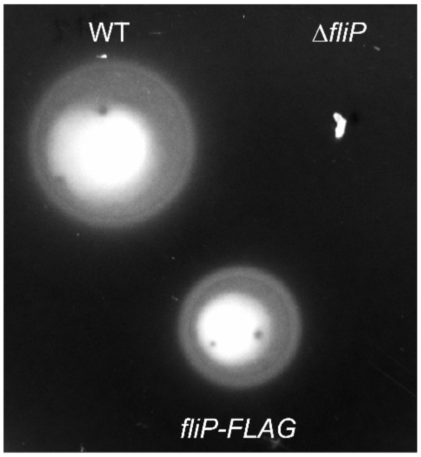

**Supplementary Fig. 2: Characterization of swimming motility of *C. jejuni* strains producing FliP-FLAG.** (A) Representative image of the motility phenotypes of WT *C. jejuni* (top left), Δ*fliP* (top right), and *fliP-FLAG* (bottom). Strains were diluted to an $OD_{600}$ of 0.8 and then stabbed into MH motility agar containing 0.4% agar. Motility was analyzed after 30 h of incubation at 37 °C in microaerobic conditions.

**Supplementary Table 1: Bacterial strains used in this study**

| Strain | Genotype | Source/Reference |
|---|---|---|
| ***E. coli* strains** | | |
| DH5α | *E. coli supE44* Δ*lacU169* (φ80*lacZ*ΔM15) *hsdR17 recA1 endA1 gyrA96 thi-1 relA1* | Invitrogen |
| DH5α / pRK212.1 | DH5α with conjugation transfer element | 1 |
| BL21(DE3) | *E. coli fhuA2 [lon] ompT gal (λ DE3) [dcm]* Δ*hsdS λ DE3 = λ sBamHIo* Δ*EcoRI-B int::(lacI::PlacUV5::T7 gene1) i21* Δ*nin5* | New England Labs |
| ***Campylobacter jejuni* strains** | | |
| DRH212 | 81-176 *rpsL*$^{Sm}$ | 2 |
| DRH706 | 81-176 *rpsL*$^{Sm}$ *fliP::cat-rpsL* | 3 |
| DRH755 | 81-176 *rpsL*$^{Sm}$ Δ*fliR* | 3 |
| DRH964 | 81-176 *rpsL*$^{Sm}$ Δ*flhA* | 3 |
| DRH1065 | 81-176 *rpsL*$^{Sm}$ Δ*fliP* | 3 |
| DRH8073 | 81-176 *rpsL*$^{Sm}$ *fliO::cat-rpsL* | This study |
| DRH8113 | 81-176 *rpsL*$^{Sm}$ Δ*fliO* | This study |
| DAR101 | 81-176 *rpsL*$^{Sm}$ Δ*fliQ* | 4 |
| SMS508 | 81-176 *rpsL*$^{Sm}$ *fliQ::cat-rpsL* | 4 |
| SNJ471 | 81-176 *rpsL*$^{Sm}$ Δ*flhB* | 5 |
| ALD962 | 81-176 *rpsL*$^{Sm}$ Δ*fliO fliR::kan-rpsL* | This study |
| ALD963 | 81-176 *rpsL*$^{Sm}$ Δ*fliO fliP::cat-rpsL* | This study |
| ALD965 | 81-176 *rpsL*$^{Sm}$ Δ*fliO fliQ::cat-rpsL* | This study |
| ALD1050 | 81-176 *rpsL*$^{Sm}$ Δ*fliO rrsC::kan-*$P_{fliNO}$*-fliO-FLAG* | This study |
| ALD1081 | 81-176 *rpsL*$^{Sm}$ Δ*fliO rrsC::kan-*$P_{fliNO}$*-*$fliO_{\Delta AMIN}$*-FLAG* | This study |
| ALD1106 | 81-176 *rpsL*$^{Sm}$ Δ*fliO* $fliP_{P22\text{-}FLAG\text{-}T23}$ | This study |
| ALD1130 | 81-176 *rpsL*$^{Sm}$ $fliP_{P22\text{-}FLAG\text{-}T23}$ | This study |

**Supplementary Table 2: Plasmids used in this study**

| Plasmid | Genotype | Source/Reference |
|---|---|---|
| pUC19 | $Amp^R$; general cloning vector | New England Biolabs |
| pBR322 | $Amp^R$; general cloning vector | New England Biolabs |
| pILL600 | $Kan^R$; source of *kan* cassette | 6 |
| pDRH265 | pUC19 containing *cat-rpsL* | 3 |
| pDRH643 | pUC19 containing *fliR*::*cat-rpsL* | 3 |
| pDRH645 | pUC19 containing *fliP*::*cat-rpsL* | 3 |
| pDRH2455 | pUC19 containing *fliO* with 0.7 kb upstream and downstream cloned into the BamHI site | This study |
| pDRH2547 | pUC19 with DNA fragment to create an in-frame deletion of codons 2 to 263 of *fliO* cloned into the BamHI site | This study |
| pDRH2559 | pUC19 containing *fliO* with 0.7 kb of upstream and downstream sequence cloned into the BamHI site; contains T410G mutation to create an StuI site in *fliO* | This study |
| pDRH2568 | SmaI-digested *cat-rpsL* cassette cloned into the StuI site of *fliO* in pDRH2559 | This study |
| pDRH7574 | pBR322 with a 2.6 kb DNA fragment containing a portion of the coding sequence of *rrsC* through a portion of *rrlC* was cloned into the EcoRI site | This study |
| pDRH7746 | *kan* cassette from pILL600 cloned into the XbaI site of the *rrsC* locus of pDRH7574 | This study |
| pSMS469 | pUC19 containing *fliQ*::*cat-rpsL* | 4 |
| pALD1047 | *fliNO* promoter from 120 bp up to the *fliN* start codon fused to the start codon of the *fliO* coding sequence with DNA encoding a C-terminal FLAG tag cloned into the XbaI site of pDRH7746 | This study |
| pALD1059 | *fliNO* promoter from 120 bp up to the *fliN* start codon fused to the start codon of the *fliO* coding sequence lacking the predicted AMIN domain (codons 21-118) with | This study |

| | | |
|---|---|---|
| | DNA encoding a C-terminal FLAG tag cloned into the XbaI site of pDRH7746 | |
| pALD1062 | *fliP* locus with 0.7 kb of upstream and downstream sequence with DNA encoding a FLAG tag between codons 22 and 23 to encode $FliP_{P22\text{-}FLAG\text{-}T23}$ cloned into the EcoRI site of pUC19 | This study |

**Supplementary Table 3: Identification by mass spectrometry of fT3SS components isolated from total membrane fractions of WT *C. jejuni* and the Δ*fliO* mutant.**

| Gene ID | Protein Name | WT/Δ*fliO* Abundance* |
|---|---|---|
| CJJ81176_0837 | FliP | 0.79 |
| CJJ81176_0357 | FlhB | 2.26 |
| CJJ81176_0890 | FlhA | 0.94 |
| CJJ81176_0340 | FliF | 1.13 |
| CJJ81176_0341 | FliG | 1.01 |
| CJJ81176_0502 | SecG | 0.99 |

*The ratio of the abundance of each protein in WT *C. jejuni* and *C. jejuni* Δ*fliO* was calculated. FliR and FliQ were not detected by mass spectrometry. SecG is a non-flagellar membrane protein that served as a control for a protein that should be at similar levels in the *C. jejuni* strains.
